# Geographic variability of hybridization between Red-breasted and Red-naped Sapsuckers

**DOI:** 10.1101/2022.08.01.502336

**Authors:** Libby Natola, Shawn M. Billerman, Matthew D. Carling, Sampath S. Seneviratne, Darren Irwin

## Abstract

Hybrid zones reveal the strength of reproductive isolation between populations undergoing speciation and are thus a key tool used in evolutionary biology research. Multiple replicate transects across the same hybrid zone offer further insight into the dynamics of hybridization in different environments, clarifying the role of extrinsic forces on the speciation process. Red-breasted and Red-naped Sapsuckers (*Sphyrapicus ruber* and *S. nuchalis*) have a long zone of contact over approximately 1,600 km from central British Columbia, Canada to central California, USA. We compared Genotyping-by-Sequencing data from three independent sapsucker hybrid zone transects to compare hybridization dynamics between the same species under variable geoclimatic conditions. We then generated geographic clines of the genomic data to compare hybrid zone widths and used Random Forest models and linear regression to assess the relationship between climate and sapsucker ancestry along each transect. Our results show variation in symmetry and directionality of back crossing, patterns often indicative of moving hybrid zones. We note variable cline widths among transects, indicating differences in the selection maintaining hybrid zone dynamics. Furthermore, Random Forest models identified different variables in close association with sapsucker ancestry across each transect. These results indicate a lack of repeatability across replicate transects and a strong influence of the local environment on hybrid zone dynamics.

## Introduction

A major goal of evolutionary biology is to determine the forces that drive the formation of new species from a shared common ancestor. Speciation occurs when some barrier between populations forms, i.e., they are no longer able to successfully interbreed. There are many ways environments may facilitate reproductive isolation between species, for example, specialization of microhabitat (*Dicurus*, Mayr 1947), soil chemistry (*Hieracium*, Turesson 1922), foraging habitat (*Gasterosteus*, Schluter & McPhail 1992), or parasite-host relationships (*Rhagoletis*, Bush 1966). Whereas we have learned much about the role of the environment from studies using experimental methods to manipulate environmental conditions for speciating groups (*Drosophila*, Matute et al. 2009; *Saccharomyces*, Dettman et al. 2007), it is difficult to study in empirical systems. Hybrid zones, in which two or more incompletely isolated forms come into contact and interbreed, serve as a great opportunity to study natural speciation in progress. However, hybrid zones are relatively rare and usually geographically small (McCarthy 2006), so it is difficult to observe species’ responses to different environmental factors. Furthermore, natural environments are stochastic and variable, clouding the associations between isolation and any one environmental barrier.

Replicate transect analysis, wherein one or more hybrid zones between two species are compared across independent transects that are geographically well separated, is a powerful tool to identify environmental features associated with reproductive barriers (Arntzen et al 2017, Culumber et al 2014, Zielinski et al 2019). For example, variable hybrid zones among spotted and collared towhees (*Pipilo maculatus* and *P. ocai*) were linked to habitat differences between the zones (Kingston et al 2014). Replicate transect analysis was also used to demonstrate lack of repeatability in swordtail fish hybridization (*Xiphophorus birchmanii* and *X. malinche*, Culumber et al. 2014) and in sculpin hybridization (*Cottus perifretum* and *C. rhenanus*, Nolte et al 2009). Replicate hybrid zones can also lend insights to hybrid zone ages and movement (Zielinski et al 2019, Arntzen et al 2017).

Hybrid zones, particularly their widths, are useful in understanding the role of selection in population isolation. Selection for or against hybrids might primarily be related to intrinsic isolation (incompatibilities independent of habitat) or to extrinsic (environment-influenced) isolation. The former would lead to similar widths across replicate hybrid zones, whereas the latter could lead to width differences if there is variation in environmental conditions between different replicate transects. Widths, and other hybrid zone parameters, can be estimated using cline theory, which tracks transitions of allele or phenotype frequencies from one species range to another across a hybrid zone (Barton & Hewitt 1985). In a standard tension zone, the width and center of a cline depend on dispersal of parental forms into the hybrid zone balanced by selection against hybrids. In natural replicated hybrid zones, selective pressures could vary due to different environments, causing different cline widths, locations, and asymmetries. Comparing differences in cline structure across different hybrid zones can therefore help us understand the isolating barriers influencing reproductive isolation. In toads *Bufo bufo* and *B. spinosus*, replicate hybrid zone transects have variable cline structure and show different numbers of barrier loci indicating variable strengths of selection among replicate transects (van Riemsdijk et al. 2020).

Red-breasted and Red-naped Sapsuckers (*Sphyrapicus ruber* and *S. nuchalis*) are widespread woodpeckers in western North America which hybridize along multiple long zones of geographical overlap, providing an excellent system to study how replicate hybrid zone transects can help us to better understand the evolution of reproductive isolation. Across one transect through the hybrid zone in central British Columbia (BC) and across another 800 km transect to the south on the California/Oregon border, hybridization (the creation of F1, F2, and advanced generation hybrids) and introgression (the repeated backcrossing of hybrids with parental species to create advanced generation backcrosses) are somewhat frequent (Billerman et al., 2016; Seneviratne et al., 2016; Trombino, 1998). However, historical records of sapsuckers within a hybrid zone in EC Manning Provincial Park in southern BC show little incidence of hybridization (Howell, 1953). These observations suggested that studying replicate hybrid zones in Red-breasted and Red-naped Sapsuckers could provide important insights into the barriers to geneflow and resulting speciation in these two species. However, the different hybrid zone transects were studied by different researchers using different metrics, so a single integrated methodology is necessary to compare the zones more directly. All three regions have been described as tension zones (Barton & Hewitt 1985, Johnson & Johnson 1985, McCarthy 2006), and so are excellent candidates for cline analysis.

In this study we set out to answer two main questions. First, to what degree do genetic cline widths and hybridization rates (i.e., proportions of population that are admixed) differ between different transects? High variability might implicate differences in the role of local habitat transitions, whereas close similarity would point to the hybrid zone dynamics being mostly independent of habitat differences (Barton & Hewitt 1989, Orive & Barton 2002). Second, are the hybrid zone widths and amount of hybridization related to the steepness of the environmental transition? We hypothesized that environmental conditions contribute to variation in hybridization and that these are likely tied to habitat transition strength. We approached this study by comparing clines from genotyping-by-sequencing data of sapsuckers at three hybrid zones: a Northern transect in central BC (Seneviratne et al. 2016), a Central transect in southern BC (this study), and a Southern transect in Oregon and California (Billerman et al. 2016). Here, we compare hybridization rates, cline structure, and landscape/climatic data to describe differences across different latitudinal transects.

## Methods

### Sample collection

Sampling was conducted in three widely separated hybrid zone transects (Figure 1). In the Northern Transect in central BC, Seneviratne et al. (2016) sampled blood from 133 sapsuckers across 556 km (−126.9217 °W, 52.4808 °N to −119.455 °W, 50.6364 °W, Figure 1). In the Central Transect in southern BC, we sampled blood from 46 sapsuckers across a 129 km transect (−120.13 °W, 49.39612 °N to −121.8936 °W, 49.23745 °N) between April 25 and July 15 in 2016, 2018, and 2020. In the Southern Transect in California and Oregon, Billerman et al. (2019) collected and vouchered 136 samples as specimens across 645 km (−117.2634° W, 45.61316° N to −123.5681° W, 42.01527° N). In the Northern and Central transects, sapsuckers were trapped using either a dip net at the nest or mist nets, decoys, and audio lures. For all BC birds, we collected ~50 μL of blood stored in Queen’s lysis buffer, and banded each individual with an aluminum band issued by the Canada Bird Banding Office to prevent resampling.

**Figure 1.**
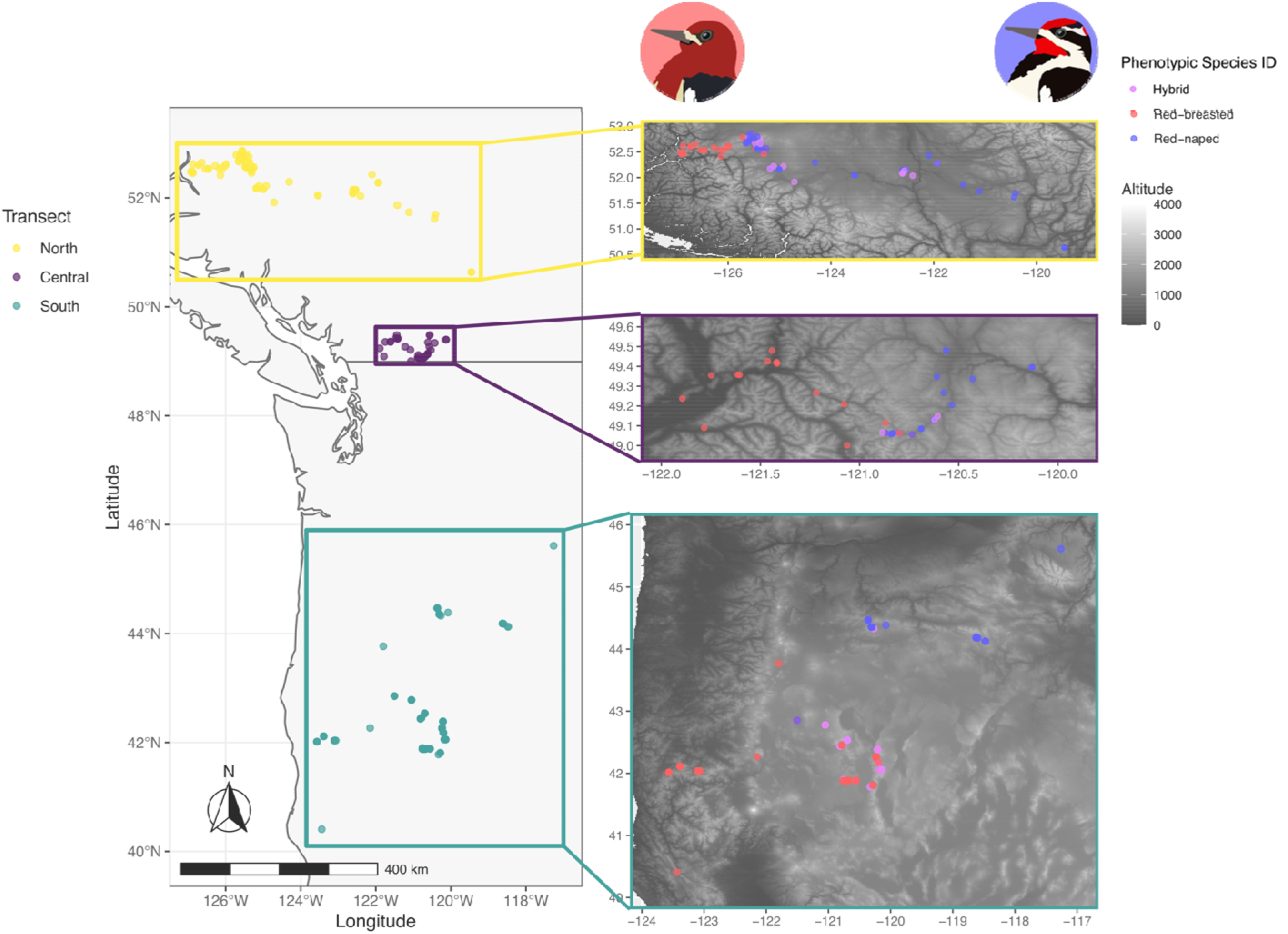
Sample distribution for the Northern (yellow), Central (purple) and Southern (teal) transects. Dots indicate sampling localities for birds phenotypically identified as Red-breasted (red), Red-naped (blue) and hybrid (violet) Sapsuckers.

### DNA extraction

DNA was extracted from samples from the Northern and Central Transects using a standard phenol-chloroform extraction. For samples from the Southern Transect, DNA was extracted by Billerman et al. (2019) using a DNeasy tissue kit (Qiagen).

### GBS protocol

For all three datasets, we followed the genotyping-by-sequencing (GBS) protocol detailed by Elshire et al. (2011) and Alcaide et al. (2014) using the enzyme *PstI*. Slight deviations from this protocol for the Southern and Northern Transects are detailed in Billerman et al. (2019) and Seneviratne et al. (2016), respectively. For the Central Transect, we followed Seneviratne et al. (2016) with the following changes. We incubated the digestion with 1 μL *PstI* (20,000 U/μL), 2 μL 10X buffer, 6μL (0.4 ng/μL) of common adaptor, 6μL (0.4 ng/μL) of barcode), and 5 μL (20ng/μL) of template DNA at 37°C for 2 hours. We ran the incubation for ligation at 22°C for 1 hour then 65°C for 10 minutes. For the PCR step, we included 5μL of 5x Phusion buffer, 0.5 μL (10mM) dNTPs, 1.25 μL (10 mM) forward and reverse primers, 12.75 μL UltraPure water, and 0.25 μL (5 U/μL) of PhusionTaq. We ran the reaction protocol as follows: 98°C for 30 seconds and 18 cycles of: 98°C for 10sec, 65°C for 30sec and 72°C for 30 sec. This was followed by an extension of 72°C for 5min and 4°C. We then selected 400-500 bp fragments. We sent 100 ng to Genome Quebec for paired-end 150bp sequencing on the Illumina HiSeq4000 or the NovaSeqSP 6000 (one library each) following Genome Quebec’s discontinuation of the HiSeq4000 platform (Illumina, San Diego, CA). The NovaSeq sequences the same reads as the HiSeq but at greater read depths, which did not affect our analyses.

### Read Processing

We ran the Tassel v 3.0.174 pipeline (Bradbury et al. 2007) to process the raw reads from all three datasets to accord with the methods used in Billerman et al. (2019). These reads were assembled *de novo* and not mapped to a reference assembly. We used the default Tassel settings with few exceptions, including a minimum tag count of 3, an error rate of 0.01, and minor allele frequency between 0.01 and 0.5. After processing reads through the Tassel pipeline, we retained 99 individuals and 106,344 SNPs for the Northern Transect, 46 individuals and 191,870 SNPs for the Central Transect, and 119 individuals and 127,599 SNPs for the Southern Transect. We filtered out SNPs missing in more than 30% of individuals and checked each dataset for relatedness among samples in VCFtools (Danecek et al. 2011). We randomly removed one individual per pair for all combinations with relatedness > 0.07 (the point at which related individuals stopped predominating PCA axes). We also removed individuals with more than 60% missing data, then re-filtered for SNPs missing in more than 30% of birds to eliminate reads that had dropped below the quality threshold when individuals were removed. After filtering, Northern had 67,092 SNPs and 76 individuals, Central included 149,660 SNPs from 43 birds, and Southern consisted of 29,236 SNPs from 108 individuals. The disparity in SNPs recovered across transects is primarily due to advances in technology (HiSeq 2000, HiSeq 4000, and NovaSeqSP 6000) and resulting differences in read depth and missing data levels, rather than a difference in genetic diversity.

### ADMIXTURE analysis

To generate PCAs from genomic data, we used the filtered, unrelated samples in each dataset separately and generated plots using methods from Irwin et al (2018). We used ADMIXTURE v. 1.3 (Alexander & Lange 2011) to estimate each sample’s ancestry in terms of the estimated proportion (Q-value) of the genome belonging to each of a given number (K) of ancestral groups. We first pruned data for LD in Plink (Purcell et al. 2007) with a window size of 50 SNPs and an r^2^ threshold of 0.1, as recommended by the ADMIXTURE documentation. The ADMIXTURE dataset pruned for LD included 27,323 SNPs for the Northern transect, 32,136 SNPs for the Central, and 13,889 SNPs for the Southern. With these pruned data, we ran ADMIXTURE for K = 1-6, computing five-fold cross-validation error and 2000 bootstrap replicates. We assigned samples as allopatric if they were outside the area where ADMIXTURE K=2 Q-values were between 0.05 and 0.95. We then used these allopatric populations to calculate genetic differentiation across each transect using Weir and Cockerham’s (1984) whole-genome weighted F_ST_.

### Geographic clines

We were also interested in the widths of the geographic clines across the three hybrid zones. To generate a one-dimensional geographic axis from the sample locations and account for differences in geographical transect orientation, we created isoclines for each transect similar to that described by Wang et al. (2019). Using Locally Estimated Scatterplot Smoothing (LOESS) in R (R Core Team), we fitted a line representing the ancestry coefficient = 0.5 based on the Q value calculated for each point in ADMIXTURE (see above), then measured each point’s geographic distance to this line as its distance to the hybrid zone (Supplement Figure 1). With this distance value as the cline x-axis, we plotted the Q-value for each sample on the cline y-axis. We first visualized the geographic extent of sympatry (overlap along transect in pure birds, Q < 0.05 or Q > 0.95) and hybridization (space along transect where admixed birds occur, 0.05 ≤ Q ≤ 0.95) in each transect. Using the R package hzar (Derryberry 2013), we plotted each transect independently and ran each of the 16 possible cline models (every combination of scaling free, fixed, or none; exponential tails both, left, right, mirrored, or none; and a null model) with a burn-in period of 100,000 and 500,000 Monte Carlo iterations. We selected the best fit model for each transect according to the lowest AICc score and include clines from those models here. We used the function hzar.getLLCutParam to calculate the upper and lower bounds of the cline centers and widths of each transect. We plotted the clines and the 95% credible interval region (the region within the maximum and minimum values of the 95% credible subset of the cline model distribution, as determined by the MCMC trace) using the hzar.plot.fzCline tool.

### Climate relationships with hybridization

To assess the relationship between hybrid ancestry at each of the three clines and environment, we used Random Forests (RF) models (Breiman 2001, Cutler et al. 2007, Evans & Murphy 2019, Franklin 2009, A. Liaw & Wiener 2002) and regression analyses. We acquired climate data for these analyses from the WorldClim database, downloading the 19 bioclimatic variables at a resolution of 0.5 minutes of a degree for tiles 11 and 12, across which our three transects span (WorldClim 2022). We downloaded these variables using the ‘*getData*’ function in the raster package v 3.5-21 (Hijmans et al. 2022) in R v. 4.2.0 (R Core Development Team 2022). We similarly downloaded elevation data for the same region. Before using these variables in our models, we first tested the 19 bioclimatic variables for multicollinearity using the rfUtilities v 2.1-5 package (Evans & Murphy 2019) as implemented in R (R Core Development Team 2022). Using a multicollinearity threshold of 10^-7^, we removed variables for each transect analyses, leaving 9, 11, and 6 nonredundant variables for the southern, central, and northern transects, respectively. We assessed the climate variables that were important for predicting ancestry values of the two species across the hybrid zone along each transect separately using RF models, using the regression mode. Ancestry values for each individual were calculated in ADMIXTURE (see Methods: ADMIXTURE analysis), and these values were used in RF models. RF models were built using the randomForest v 4.7-1.1 (M. Liaw 2022) and rfUtilities v 2.1-5 (Evans & Murphy 2019) packages as implemented in R. Models were run using the following conditions: 1,000 trees (bootstrap iterations) and using the “mir” scale, which is the permutation-based importance measure for each predictor variable. We selected our final model by maximizing the amount of variation explained, minimizing the root mean squared error (MSE), and maximizing parsimony (minimize the number of parameters) to maximize ecological interpretability. Fit statistics for each RF model were also assessed using the ‘*rf.regression.fit*’ function, which also evaluates whether the model is overfit. After selecting our top models for each transect, we ran separate linear regression models as implemented in R (R Core Development Team 2022) to assess significance, direction, and magnitude of the influence of each variable on predicting hybrid ancestry.

### Enviroclines

To visualize the relationship between the hybrid dynamics and environment, we plotted the genomic clines together with the environmental data, hereafter referred to as enviroclines. We drew a horizontal latitudinal transect bisecting the isocline midpoint along which we plotted data from the three environmental variables included in each transect’s RF model in conjunction with our hzar clines (Supplement Figure 1). We note that these linear transects are useful in distilling complex geographic and climate information in a simplistic visualization, but may show data from regions uninhabited by sapsuckers, particularly in the patchier landscape of the Southern transect.

## Results

### Genomic differentiation

Each of the hybrid zone transects shows clear genomic differentiation between Redbreasted and Red-naped Sapsuckers, but there are strong differences in the proportion and genomic ancestry of phenotypic hybrids as well as the structure of environmental change. In the Central Transect, there was only one intermediate individual which was located precisely between the two parental species clusters, whereas in the Northern Transect phenotypic hybrids tended to group genetically with Red-naped Sapsuckers and in the Southern Transect phenotypic hybrids were mostly positioned with the Red-breasted Sapsuckers (Figure 2). As these were visually identified as hybrids by different researchers, we inspected ancestry data to infer whether birds with admixed genotypes cluster with different parental groups in different hybrid zone transects or if field researchers had different phenotypic criteria for classifying intermediate birds.

**Figure 2.**
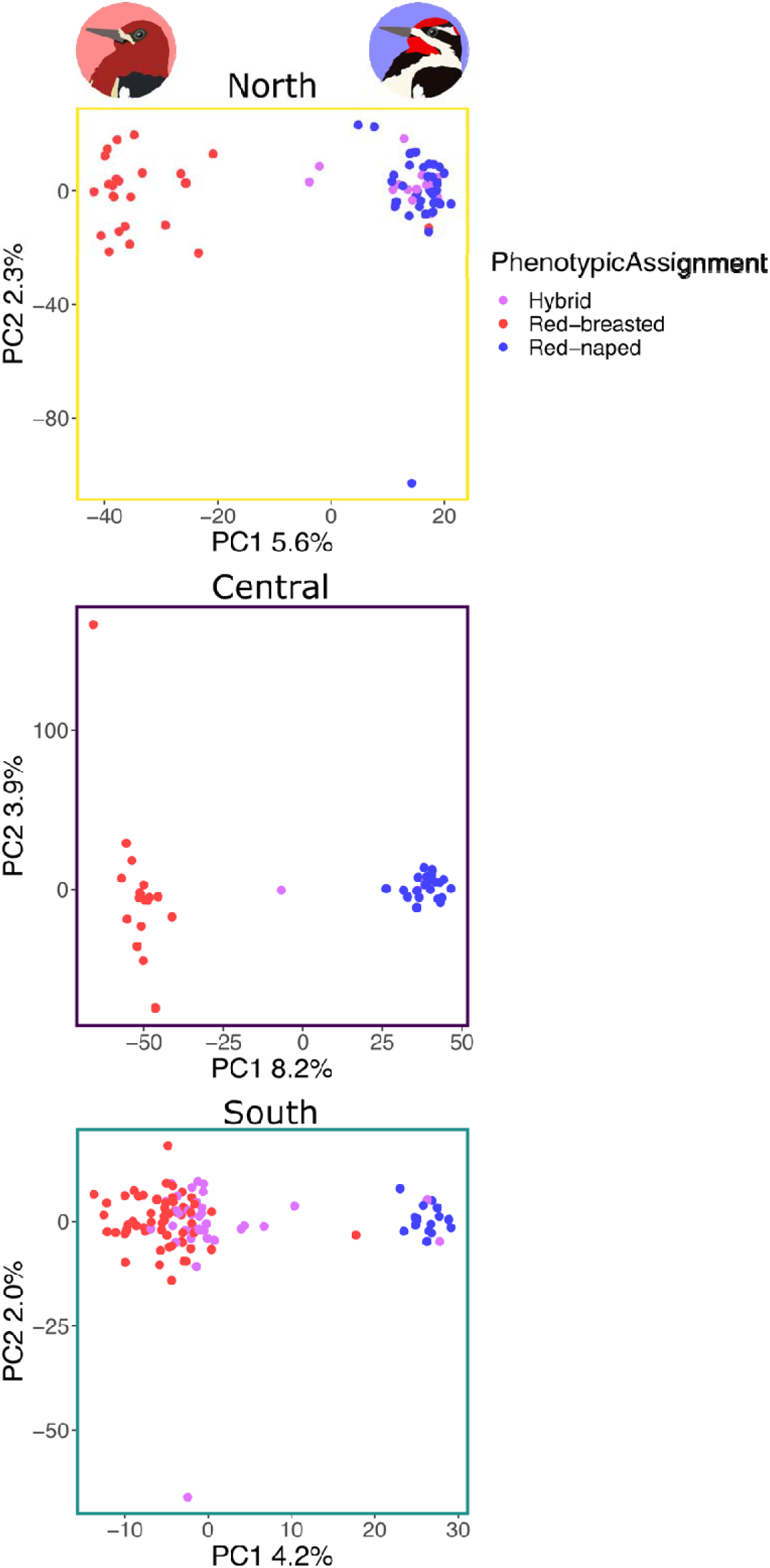
Principal Component Analyses of Northern (top, yellow), Central (middle, purple), and Southern (bottom, teal) transect genomic data. Dots colored according to phenotypic species assignment based on plumage for hybrid (violet), Red-breasted (red), and Red-naped Sapsuckers (blue).

### ADMIXTURE analysis

We used K=2 in ADMIXTURE, because of the *a priori* expectation that these zones are meeting places of two species. While K=3 might in some cases show hybrids as an identifiable entity, K=2 had lower cross-validation error, and allowed us to estimate ancestry contributions from each parental species in admixed birds. This resolved the species well in a histogram. We accepted the K=2 model for each transect based on these visual plot examinations (Figure 3). Like the PCAs, the Central ADMIXTURE plot showed one intermediate individual and few backcrosses, whereas the Northern plot showed admixed individuals tend to have more Red-naped ancestry, and the Southern plot showed admixed individuals tend to have more Redbreasted ancestry. In the Northern Transect 22 out of 76 (28.9%) birds were admixed (0.05 < Q < 0.95), in the Central Transect 5 of 43 (11.6%) birds were admixed, and in the Southern Transect 48 of 109 (44.0%) birds were admixed. A χ^2^ test of these proportions shows the transects have significantly different proportions of admixed individuals (χ=15.863, *p* < 0.001). Further χ^2^ tests of each species pair show marginal difference between the Northern and Central transects (χ=3.76, *p* = 0.052) and Northern and Southern transects (χ=3.91, *p* = 0.048), and a large difference in the hybrid frequencies of Central and Southern transects (χ=13.135, *p* < 0.001). Allopatric Red-breasted and Red-naped Sapsucker populations in Northern, Central, and Southern transects had quite similar pairwise F_ST_ values of 0.062, 0.063, and 0.056, respectively.

**Figure 3.**
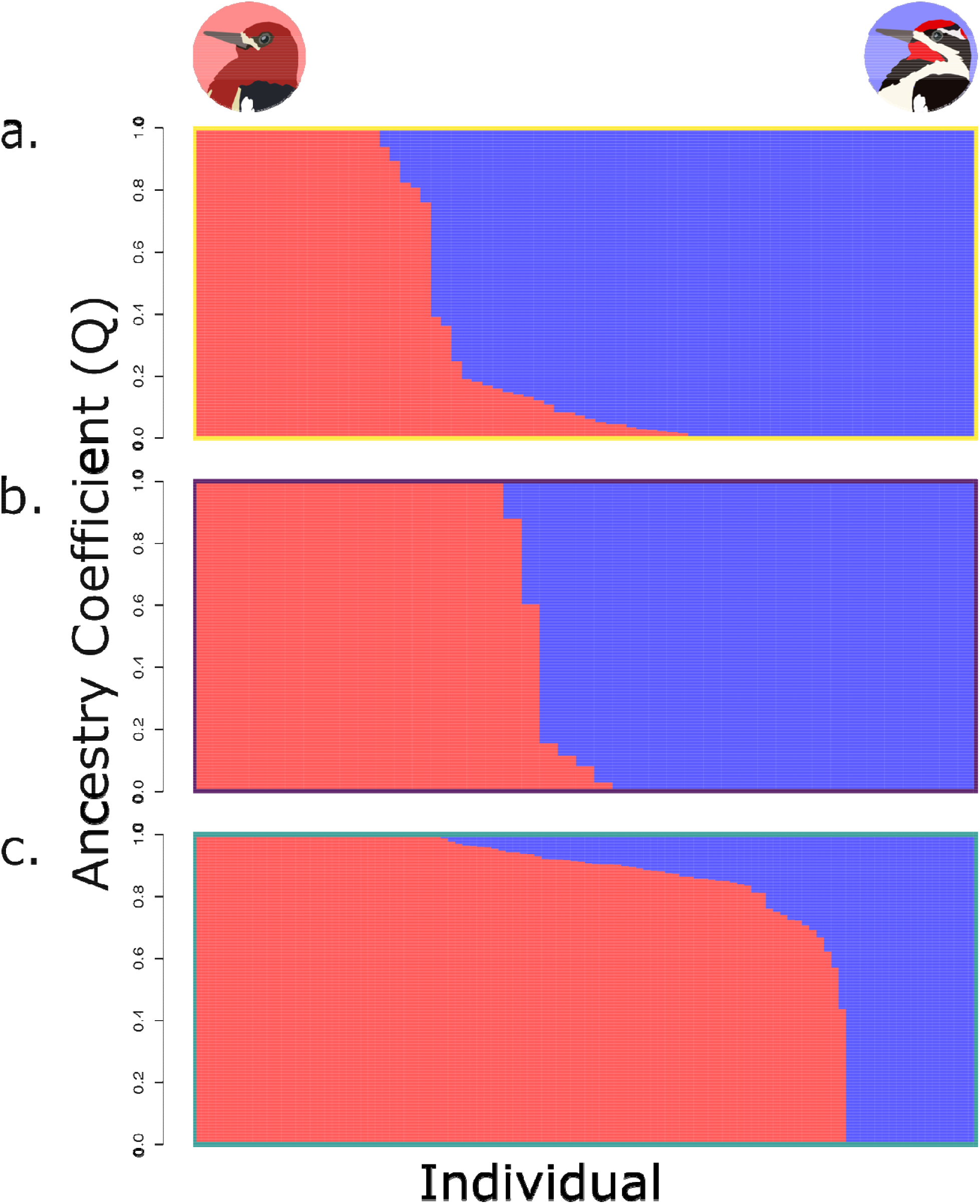
ADMIXTURE results with K=2 for Northern (a), Central (b), and Southern (c) transects. Red indicates Red-breasted Sapsucker ancestry and blue indicates Red-naped Sapsucker ancestry.

### Geographic clines

We found sympatry occurred across 14.3, 5.0, and 53.9 km (measured as distance from the isocline) and hybridization across 260.4, 13.3, and 212.9 km for the Northern, Central, and Southern transects, respectively (Supplement Figure 2). In our hzar analysis, we chose the best fit model for each transect cline based on the AICc output. For Central and Southern, the best fit was no scaling and no exponential tail, in the Northern it was no scaling and right skewed exponential tail. Our hzar analyses identified sigmoidal clines in each of the transects (Figure 4). The Central Transect was estimated at 20.0 km wide (CI: 10.5-47.7 km) and centered 0.09 km west of the isocline (CI: 8.2 km east −5.6 km west), the Northern Transect measured 27.6 km across (CI: 10.18-63.1 km) and centered 0.2 km east of the isocline (CI: 5.6 km east-13.7 km west), and the Southern Transect stretched across 138.0 km (CI: 79.3-228.6 km) centered around a point 41.1 km east of the isocline (CI: 69.7 km east-2.4 km west). The confidence intervals of cline width in the Northern transect overlapped with those of Central, but the lack of overlap between confidence intervals for Southern and either Northern or Central indicates these cline widths are different (Supplement Figure 3).

**Figure 4.**
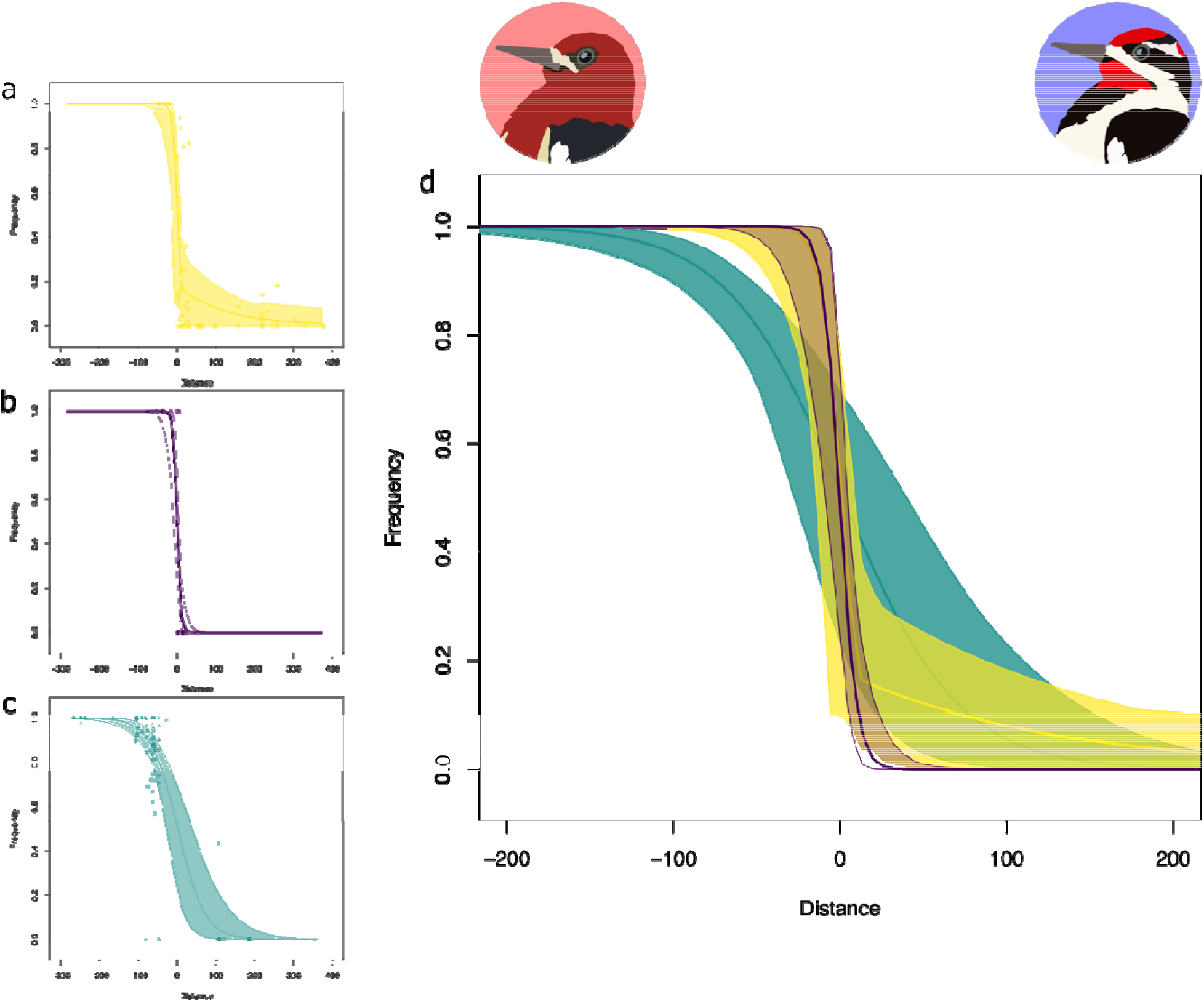
Centered genomic clines for Northern (a, yellow), Central (b, purple), and Southern (c, teal) transects. Centered sample clines (d) are plotted together. Distance on the x-axis indicates distance of each sample from the cline center estimated by hzar. Frequency on the y-axis indicates the proportion of each sample’s genome attributed to Red-breasted Sapsucker ancestry in the program ADMIXTURE.

### Climate Relationships with Hybridization

We assessed the relationship between climate and hybrid ancestry along each of the three transects separately using Random Forests models and linear regression. After removing highly correlated variables (see Methods), RF models were run with 9, 6, and 8 variables for the Southern, Central, and Northern transects, respectively. The top-ranked RF model for the Southern transect had 3 variables, including Precipitation of the Driest Month, Precipitation of the Warmest Quarter, and Mean Temperature of the Wettest Quarter (Table 1), and explained 73.58% of variation in ancestry. After testing fit of the model, we found that R^2^ was 0.734, the model had a root-mean-squared of 0.182, an accuracy of 0.922, and was not overfit. After running a linear regression, we found that Precipitation of the Driest Month had a significant negative effect on ancestry, Precipitation of the Warmest Quarter also had a negative effect on ancestry, and Mean Temperature of the Wettest Quarter had a positive effect on ancestry (Table 1).

**Table 1.** Bioclim variables used in Random Forests models. Dashes indicate that a variable was removed before the model was run due to multicollinearity. Numbers indicate the ranked importance of each variable for Random Forests models. Gray shading indicates the variable was not included in the top-ranked model.

|  | Northern | Central | Southern |
| --- | --- | --- | --- |
| Annual Mean Temperature | 5 | 4 | 4 |
| Mean Diurnal Range | 3 | 1 | 6 |
| Isothermality | 6 | -- | 9 |
| Temperature Seasonality | 1 | -- | -- |
| Maximum Temperature of the Warmest Month | -- | -- | -- |
| Minimum Temperature of the Coldest Month | -- | -- | -- |
| Temperature Annual Range | -- | -- | -- |
| Mean Temperature of Wettest Quarter | 7 | 5 | 3 |
| Mean Temperature of Driest Quarter | 4 | 6 | -- |
| Mean Temperature of Warmest Quarter | -- | -- | -- |
| Mean Temperature of Coldest Quarter | -- | -- | -- |
| Annual Precipitation | 2 | 2 | 8 |
| Precipitation of Wettest Month | -- | -- | 7 |
| Precipitation of Driest Month | -- | -- | 1 |
| Precipitation Seasonality | 8 | 3 | -- |
| Precipitation of Wettest Quarter | -- | -- | -- |
| Precipitation of Driest Quarter | -- | -- | -- |
| Precipitation of Warmest Quarter | -- | -- | 2 |
| Precipitation of Coldest Quarter | -- | -- | -- |
| Elevation | -- | -- | 5 |

The top-ranked RF model for the Central transect had 4 variables, including Mean Diurnal Range, Annual Precipitation, Precipitation Seasonality, and Annual Mean Temperature (Table 1), and explained 63.89% of variation in ancestry. After testing fit of the model, we found that R^2^ was 0.635, the model had a root-mean-squared of 0.290, an accuracy of 0.972, and was not overfit. After running a linear regression, we found that Mean Diurnal Range had a significant negative effect on hybrid ancestry, and Precipitation Seasonality had a significant positive effect on hybrid ancestry (Supplemental Table 1). Annual Precipitation also had a negative effect on hybrid ancestry, while Annual Mean Temperature had a positive effect on ancestry.

The top-ranked RF model for the Northern transect had only 2 variables, including Temperature Seasonality and Annual Precipitation (Table 1), and explained 75.97% of variation in ancestry. After testing fit of the model, we found that R^2^ was 0.754, the model had a rootmean-squared of 0.210, an accuracy of 0.248, and was not overfit. After running a linear regression, we found that Annual Precipitation had a significant positive effect on hybrid ancestry, and Temperature Seasonality had a negative effect on ancestry. In each transect, Redbreasted and Red-naped Sapsuckers (i.e., high and low Q values) tended to be separated by regions of high elevation (~1600 m) (Figure 5).

**Figure 5.**
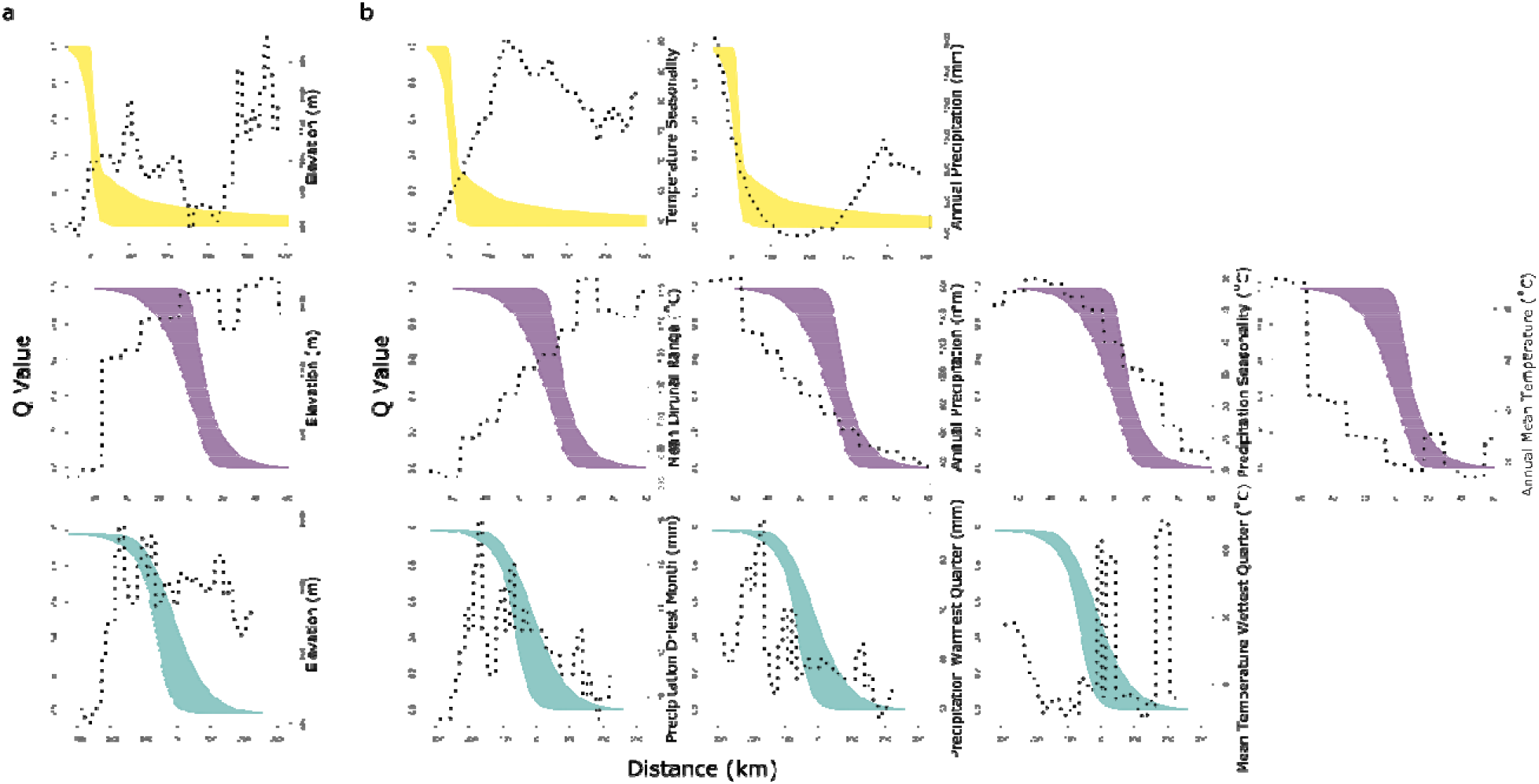
Genomic clines shown in shaded regions plotted against environmental shifts (dotted lines) of (a), elevation, and (b), the variables identified by the top RF model for the Northern (Temperature Seasonality and Annual Precipitation in mm), Central (Mean Diurnal Range in °C, Annual Precipitation in mm, Precipitation Seasonality in °C, and Annual Mean Temperature in °C), and Southern (Precipitation of Driest Month in mm, Precipitation of Warmest Quarter in mm, and Mean Temperature of Wettest Quarter in °C) transects.

## Discussion

Our results show that while some factors remain consistent among replicate Redbreasted x Red-naped Sapsucker transects, the direction and extent of hybridization vary. Genomic differentiation of non-admixed birds is comparable among transects, as demonstrated by F_ST_, and species from each transect formed two distinct and well-supported clusters (Figures 2, 3). However, the proportion of birds with admixed genomes varied among transects (28.9% Northern, 11.6% Central, 44% Southern), and the proportion and directionality of backcrossing varied from transect to transect as well. Taken together, these data demonstrate rates or directionality of hybridization and introgression are independent of the two species’ genomic background, and likely affected by extrinsic factors. Cline widths were narrower in the Northern and Central transects than they were in the Southern transect (Figure 4). These genomic transitions seem to track elevational changes and variation in annual precipitation and temperature trends, suggesting a strong role of geography and climate in the maintenance of Red-breasted and Red-naped Sapsucker hybrid zones. We are rarely afforded the opportunity to replicate empirical hybridization studies, so this contributes to the relatively small pool of data concerning the repeatability and predictability of natural introgressive systems. It also helps us put into perspective the generalizability of typically un-replicated empirical hybrid zone research.

Some research on replicate hybrid zones has found parallelisms in hybridization patterns among different transects—examples include Yellow-rumped Warblers (*Setophaga coronata coronata* and *S. c. auduboni*, Brelsford & Irwin 2009) and sunflowers (*Helianthus annuus* and *H. petiolaris*, Rieseberg et al. 1999). However, it appears that inconsistencies are a more common phenomenon. Arntzen et al. (2017) and Riemsdijk (2020) found variability in introgression of common and spiny toads (*Bufo bufo* and *B. spinosus*), which they attributed to hybrid zone movement. Swordtail fishes have inconsistencies across seven different tributaries (*Xiphophorus malinche* and *X. birchmanni*, Culumber et al. 2011) attributed to migration, selection, and parental frequency differences. In house mouse subspecies, (*Mus musculus musculus* and *M. m. domesticus*, Janoušek et al. 2012), overall patterns of introgression were not congruent, although some parallelisms were found in loci thought to be involved in intrinsic incompatibilities and epistasis. Mandeville et al. (2015) conclude that the fitness and ecological compatibility of hybrids are central to understanding inconsistent hybridization outcomes.

### Asymmetrical backcrossing

Asymmetrical introgression is not unusual. The “tails” of trailing ancestry that we document across latitudinal clines in sapsuckers have been explained in other species as being caused by moving hybrid zones. As one species’ breeding range shifts geographically, a tail of admixed genotypes remains where the hybrid zone used to occur, while there is little admixture at the leading edge of the zone where there has been a short history of secondary contact between the individuals in this area (Arntzen et al. 2017, Wielstra et al. 2017). This concept is well supported by both empirical examples (Arntzen et al. 2017, Barton & Hewitt 1981, Buggs 2007, Wielstra et al. 2017) and theory (Currat et al. 2008). Buggs (2007) described many causes for asymmetrical introgression caused by hybrid zone movement such as differences in fitness of hybrids from different parental crossings, fitness of parental species, attractiveness of parental species, allele dominance, species density, or climate change.

The asymmetrical nature of backcrossing towards Red-naped Sapsuckers in the Northern transect might thus be caused by a western movement of this hybrid zone. In 1976 Scott et al. (1976) demonstrated a northward shift of this hybrid zone based on Howell’s 1952 descriptions (Howell 1952). While they did not describe the full extent of the hybrid zone or longitudinal changes in its location, we can deduce that this zone is not static. Furthermore, approximately 275 km north of our Northern transect where Red-breasted Sapsuckers hybridize with Yellow-bellied Sapsuckers (*S. varius*), and that hybrid zone is moving west, which Seneviratne et al. (2012) suggested is caused by an expansion of Yellow-bellied Sapsuckers outcompeting the Red-breasted. It is plausible that the Red-breasted range (and consequently the hybrid zone) is also tracking westward just a few hundred km south, though whether that might result from Red-breasted range retraction or Red-naped range expansion is unclear. Changes in the forest habitats from old growth forests to monoculture forest plantations and changes in the climate could have facilitated the dry-adapted Yellow-bellied Sapsucker range to expand in the north (Seneviratne et al. 2012).

The only transect with no evidence of hybrid zone movement is the Central transect. In 1952, Howell placed the center of the zone around Allison Pass (49.1167, −120.8667) (Howell 1952). Sixty-eight years later, Allison Pass is well within the confidence interval of, and only 1.89 km from the estimated cline center. Furthermore, this region is also the only transect with symmetrical backcrossing (Figure 3).

Billerman et al. (2016) showed that the hybrid zone in the Southern Transect is shifting to the east, which is concordant with the backcrossing skew towards Red-breasted Sapsuckers (Figure 3). Arntzen et al. (2017) suggested that most incipient hybrid zones shift and don’t stabilize until they have reached an equilibrium, which is dependent upon environmental stability. Billerman et al. (2016) demonstrated a clear link between the eastward movement of the South transect and changing climate.

The inconsistencies in these zone movements across different transects can identify several of Buggs’ (2007) aforementioned hypotheses as unlikely, such as intrinsic fitness differences between parental species or different direction crosses, or dominance effects, which would show repeated uni-directional shifts. It is possible, but in our opinion unlikely, that hybrids in the Northern transect maintain an innate preference to breed with Red-naped Sapsuckers, resulting in more back-crossing in this direction and a higher proportion of Red-naped ancestry within the transect, while the South transect hybrids inherently prefer Red-breasted sapsuckers with the opposite effects. Billerman et al. (2019) did not find any explicit evidence for this preference in the South transect. While there is evidence that environmental factors (Robinson et al. 2012), hybrid zone age (Curry et al. 2016), and individual hybrid class (Fernando et al. 2016, Huynh et al. 2011) can influence mate preference and mate choice, we do not know of an example of such contradictory mate choice preferences among individuals from different populations of the same species. Northern and southern Red-breasted sapsuckers do belong to separate sub-species, which may potentially harbor opposite mating preferences, but we doubt this is likely driving the inconsistent hybrid zone patterns. Intrinsic incompatibilities between Red-breasted and Red-naped Sapsuckers likely exist, as Trombino (1998) showed hybrids are less likely to return to their breeding ranges in subsequent years and thus may have lower adult survival rates. However, in light of the directional differences in hybrid zone asymmetry, and given that there are differences in environment and selective pressures across the separate transects, the explanation for variable hybrid zone directionality that makes the most sense is that extrinsic differences in fitness of parental or hybrid sapsuckers have caused hybrid zones to move in different directions. This underscores the role of the environment in hybrid zone stability and movement. However, Buggs (2007) and Curry (2016) both concede alternate explanations for these patterns are possible, and rigorous simulations of moving hybrid zones would be necessary to ascertain the likelihood of these ancestry tails.

### Environmental influence on hybrid zone location and width

Given that sapsucker hybrid zones are prone to movement, the question arises: where do their hybrid zones tend to occur or stabilize? One variable shown to interact consistently with genotype across the enviroclines of all three transects was elevation. Each transect showed regions of high elevation (> 1,600 m) separated parental types of each species. Mountain ranges may be difficult to cross, forming a physical barrier to dispersal, and species may be ill-adapted to harsh alpine habitats (cold, low oxygen concentration, lack of trees). Though for migratory birds, it is probably more biologically relevant that alpine conditions and factors such as rain shadow effects delineate distinct biomes to which species may be differentially adapted. The location of the hybrid zone can therefore be delimited by the location of high altitude regions of the mountain range, with species ranges separated by major ridgelines. What’s more, mountains which limit contact between parental forms may act as a bulwark against gene flow, preventing movement of alleles from crossing to either side of the range, narrowing hybrid zone widths. Our enviroclines do show this division of genotypes on either side of major ridgelines (Figure 5). Furthermore, the Northern and Central transects, which had the closest association to elevation, also show comparably narrow hybrid zone widths. However, it is worth noting that the broader hybrid zone in the Southern transect has a similarly high elevation peak to the Northern and Central transects.

Another way to infer selective forces influencing hybrid zone width may be to identify variables that covary with genotype transition across space. For example, if one climatic trait was tightly correlated with ancestry coefficients across multiple transects, we might infer a possible causal relationship between these. However, our RF models indicate that sapsucker hybrid zone dynamics are not responding to any one climatic variable. The variables identified as important in each transect are different (with the exception of Annual Precipitation, which was correlated with ancestry in both the Northern and Central zones), indicating that the variable hybrid zone transitions are associated with different climatic regimes. According to the RF models, annual patterns in temperature and precipitation explain the majority of variation in the location of sapsucker ancestry.

While precipitation and temperature are clearly implicated in sapsucker spatial distribution, each is in turn also affected by elevation, making it difficult to entirely decouple these variables. Furthermore, these three factors will all influence ecological patterns such as forest composition, which also determines species distributions. The hybrid zone occurs across a vast latitudinal gradient, and each transect exhibits considerable differences in ecology, climate, and topography. It is possible that habitat specialization and habitat transitions preclude introgression more in some transects than others. Red-breasted Sapsuckers live primarily in temperate Pacific coast climates, preferring coniferous forests dominated by western white pine (*Pinus monticola*), lodgepole pine (*Pinus contorta*), and western hemlock (*Tsuga heterophylla*) (Walters et al. 2002). Red-naped Sapsuckers live in the more arid intermountain west region in deciduous or mixed forests predominated by aspen (*Populus tremuloides*) trees (Walters et al. 2002). Hybrid zones that exist on a sharp ecotone might be narrower than those on a landscape that presents a gradual transition between parental species’ preferred habitats or wide swathes of mutually occupied habitats. Anecdotally, our narrowest transect (Central) occurs across a very steep ecotone, while our broadest (Southern) exists on a broad and patchy habitat transition. It might be imagined that if each species is differentially adapted to a climate variable, the rapid turnover of that variable in space may reduce the availability of intermediate habitat where hybrids are often found to have competitive or superior fitness with parental forms (Good et al. 2000, Hamilton et al. 2013). This steep transition in many of the variables identified by the RF models may therefore make it difficult for hybrids to find their ecological toehold, keeping hybrid zones narrow.

Barton and Hewitt’s (1989) equation describing cline widths in hybrid zones identifies high selection against hybrids and low parental dispersal into the hybrid zone as two major causes for narrow hybrid zones. Our enviroclines analysis suggests possible explanations involving both of these mechanisms. High elevation areas and associated habitat changes may reduce dispersal in and out of hybrid zones, and limited intermediate habitat caused by sharp environmental transitions may adversely affect hybrid fitness. Local adaptation is complex, and climate and habitat variables are inherently coupled. It is therefore not possible to calculate the exact effect every selective force contributes to empirical species divergence. However, such qualitative research allows us to examine hybrid zone behaviour under different environmental regimes. A limitation of our study is that we do not have the raw data files for all our datasets and are unable to map our reads to a reference genome to identify outlier loci. It would be interesting to compare the loci under selection in the different transects to shed more light on the selective forces driving the patterns described here.

## Conclusions

Our data demonstrate that though differentiation between Red-breasted and Red-naped Sapsuckers is consistent across three separate transects of their ranges, the species interactions are quite different across independent transects with different hybridization rates, skews, and cline widths. This suggests hybridization dynamics are not highly repeatable and implicates a strong effect of the environment in shaping local hybridization dynamics. Hybrid zone movement may also play a role in skewing clines, and previous research shows movement is likely mediated by climatic factors.

## Acknowledgements

K. Martin, L. Rieseberg, and D. Schluter provided key feedback on all stages of project development. A. Ali, A. Bartels, M. R. Jones, J. M. Maley, E. Nikelski, G. Pang, J. R. Saucier, K. Wang, and C. Welke assisted in field sample collection. A. Bazzicalupo, M. Boehm, T. Booker, C. Elphinstone, A. Geraldes, J. Grummer, and M. Whitlock provided feedback on analyses. This research was funded by the Natural Sciences and Engineering Research Council of Canada (RGPIN-2017-03919 and RGPAS-2017-507830 to D.I.) and the Warner and Hildegard Hesse Research Award (to L.N.). S.M.B. was supported by a National Science Foundation Graduate Research Fellowship, the American Museum of Natural History Frank M. Chapman Memorial Fund, a Sigma-Xi Grant-in-Aid of Research, the University of Wyoming Department of Zoology and Physiology, the University of Wyoming Program in Ecology, and the University of Wyoming Museum of Vertebrates. We thank T.Burg (University of Lethbridge), S. Birks and R. C. Faucett (University of Washington Burke Museum of Natural History and Culture), G. Hanke (Royal BC Museum) J. Hudon (Royal Alberta Museum), C. Stinson and I. Szabo (Beaty Biodiversity Museum), for sharing specimens and/or tissue samples. For research permits we thank the UBC Animal Care Committee, the Canadian Bird Banding Office, Environment and Climate Change Canada, BC Parks, the United States Fish and Wildlife Service, the Oregon Department of Fish and Wildlife, the Washington Department of Fish and Wildlife, the California Department of Game and Fish, and the University of Wyoming IACUC.

## Data Accessibility

All code used in these analyses is shared in a GitHub repository (https://github.com/libby-natola/Variable-hybridization-Red-naped-and-Red-breasted-Sapsuckers) which will be made public upon publication.

