## Supplement Figure 1 for "Geographic variability of hybridization between Red-breasted and Red-naped Sapsuckers"

Supplemental Table 1. Linear regression of variables identified by the top-ranked RF model for each transect. The values marked with a * indicate variables that pass significance of p < 0.05, *** indicate significance beyond p < 0.001.

| Transect | Variable | Estimate | t-value | P(>\|t\|) |
| --- | --- | --- | --- | --- |
| Northern | Intercept | 8.08E-01 | 1.198 | 0.235 |
|  | Temperature seasonality | -1.44E-04 | -1.941 | 0.056 |
|  | Mean annual precipitation | 8.98E-04 | 3.909 | 0.00021* |
| Central | Intercept | 5.984 | 2.241 | 0.031* |
|  | Mean diurnal range | -0.052 | -2.433 | 0.020* |
|  | Mean annual precipitation | -0.001 | -1.231 | 0.223 |
|  | Precipitation seasonality | 0.022 | 2.165 | 0.037* |
|  | Mean annual temperature | 0.002 | 0.382 | 0.705 |
| Southern | Intercept | 1.917 | 9.686 | < 0.0001*** |
|  | Precipitation driest month | -0.052 | -2.525 | 0.013* |
|  | Precipitation warmest quarter | -0.007 | -1.439 | 0.153 |
|  | Mean temperature warmest quarter | 0.005 | 1.485 | 0.141 |

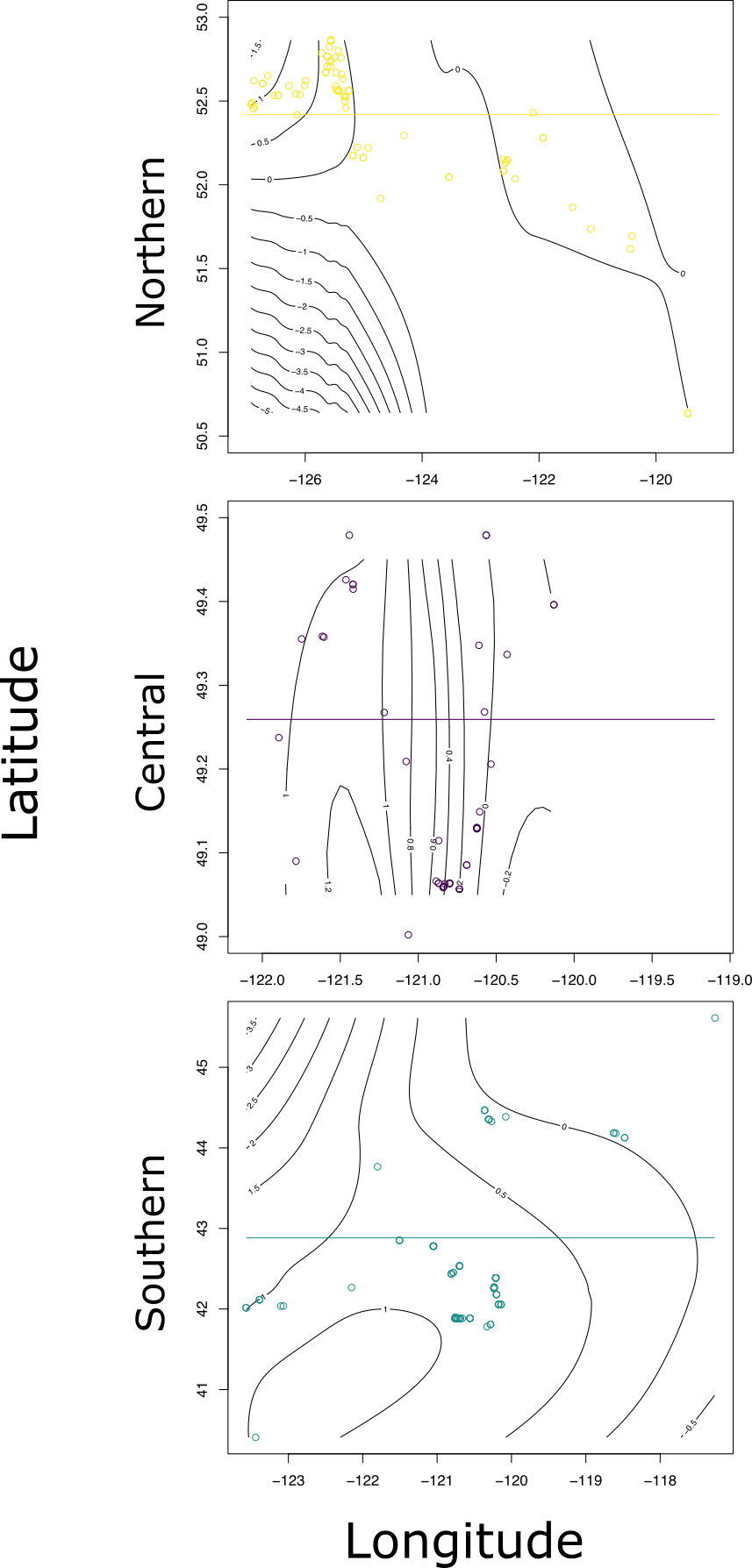

Supplement Figure 1. Isoclines of ancestry coefficients shown in black with sample locations shown as circles and the transects used for the enviroclines shown in yellow, purple, and teal for Northern, Central, and Southern transects respectively.

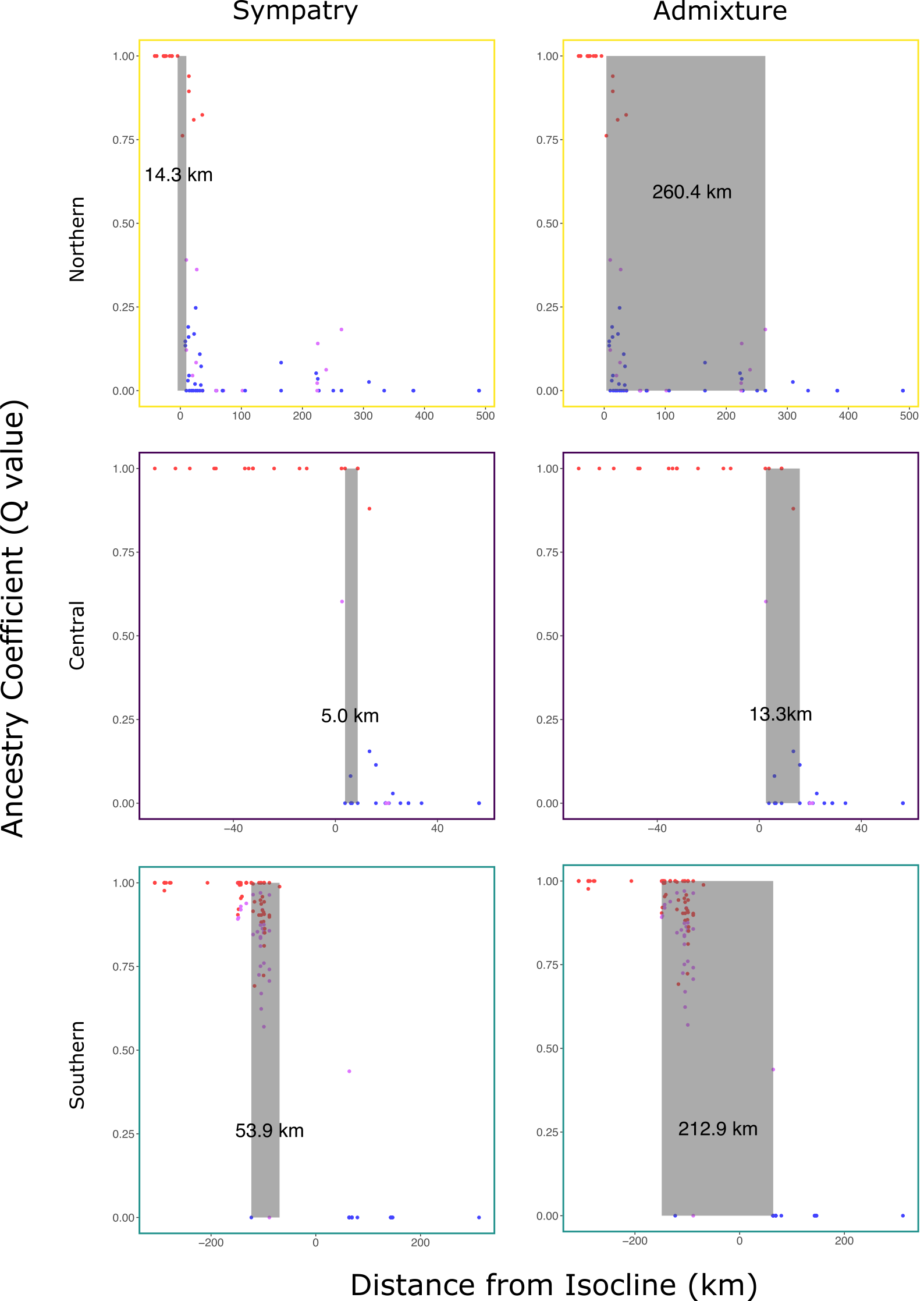

Supplement Figure 2. Samples plotted as distance from isocline by ancestry coeffiecient (Q). Shaded boxes indicate regions of sympatry (left) and admixture (right) for Northern (top, yellow), Central (middle, purple), and Southern (bottom, teal) transects. Dot colors denote phenotypic species classifications as Red-breasted (red), Red-naped (blue) or hybrid (violet) Sapsuckers.

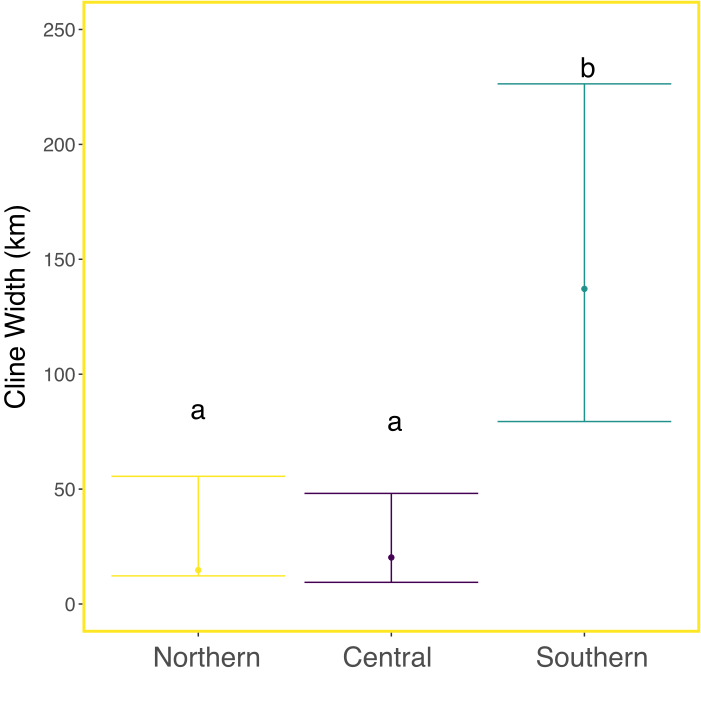

Supplement Figure 3. Estimated cline width with confidence intervals for the Northern (yellow), Central (purple), and Southern (teal) transects.
